# The Na^+^(K^+^)/H^+^ exchanger Nhx1 controls multivesicular body-vacuolar lysosome fusion

**DOI:** 10.1101/170175

**Authors:** Mahmoud Abdul Karim, Christopher Leonard Brett

## Abstract

Loss-of-function mutations in human endosomal Na^+^(K^+^)/H^+^ Exchangers (NHEs) NHE6 and NHE9 are implicated in neurological disorders including Christianson Syndrome, autism and attention deficit and hyperactivity disorder (ADHD). These mutations disrupt retention of surface receptors within neurons and glial cells by affecting their delivery to lysosomes for degradation. However, the molecular basis of how these endosomal NHEs control endocytic trafficking is unclear. Using *Saccharomyces cerevisiae* as a model, we conducted cell-free organelle fusion assays to show that transport activity of the orthologous endosomal NHE Nhx1 is important for multivesicular body (MVB)-vacuolar lysosome fusion, the last step of endocytosis required for surface protein degradation. We find that deleting Nhx1 disrupts the fusogenicity of the MVB, not vacuole, by targeting pH-sensitive machinery downstream of the Rab-GTPase Ypt7 needed for SNARE-mediated lipid bilayer merger. All contributing mechanisms are evolutionarily conserved offering new insight into the etiology of human disorders linked to loss of endosomal NHE function.

## INTRODUCTION

Found in all organisms, Na^+^(K^+^)/H^+^ Exchangers (NHEs) are secondary active ion transporters that conduct exchange of monovalent cations for hydrogen ions across membranes, and thus contribute to diverse physiology by regulating compartmental osmolytes, volume and pH (*Brett et al., 2005a*). Nhx1, an NHE in *S. cerevisiae,* resides on endosomal compartments within cells (*Nass and Rao, 1998; Kojima et al., 2012*). Originally identified as an important contributor to salt homeostasis (*Nass et al., 1997*), Nhx1 was later discovered to play a pivotal role in membrane trafficking (*Bowers et al., 2000*): Also called VPS44, NHX1 is one of ~60 *vps* (Vacuole Protein Sorting) genes originally identified in series of classic genetic screens that revealed the molecular underpinnings of endocytic trafficking (*Robinson et al., 1988; Rothman et al., 1989*). A hallmark phenotype of NHX1 knockout (*nhx1*∆) cells and other *vps* class E mutants is an aberrant, enlarged late endosomal compartment where internalized surface proteins and biosynthetic cargoes get trapped and accumulate en route to the vacuolar lysosome (*Bowers et al., 2000; Brett et al., 2005b*). Other *vps* class E mutants have since been shown to encode loss-of-function mutations in components of the ESCRT (Endosomal Sorting Complex Required for Transport) machinery responsible for sorting and packaging internalized surface receptor and transporter proteins into intralumenal vesicles at the endosome (*Henne et al., 2011*). Many rounds of ESCRT-mediated intralumenal vesicle formation produce mature multivesicular bodies (MVBs) that then fuse with lysosomes exposing protein-laden vesicles to acid hydrolases for catabolism. Because cells devoid of ESCRTs or NHX1 share many phenotypes, we originally hypothesized that deletion of NHX1 also prevented intralumenal vesicle formation. However, further studies revealed that intralumenal vesicle formation persists in *nhx1*∆ cells (*Kallay et al., 2011*), albeit less efficiently than in wild type cells (*Mitsui et al., 2011*).

Based on data from phenomics screens, a new model of Nhx1 function emerged proposing that endocytic defects observed in *nhx1*∆ cells may be caused by MVB membrane fusion defects (*Kallay et al., 2011*). Lending the greatest support to this hypothesis is the finding that Nhx1 binds Gyp6 (*Ali et al., 2004*), a Rab-GTPase Activating Protein (Rab-GAP) that inactivates two Rab-GTPases thought to drive membrane fusion (*Strom et al., 1993; Vollmer et al., 1999; Will and Gallwitz, 2001*): Ypt6, for MVB – Trans-Golgi Network (TGN) vesicle fusion (*Bensen et al., 2001; Luo and Gallwitz, 2003; Suda et al., 2013; Brunet et al., 2016*) and Ypt7, for MVB – vacuolar lysosome (or vacuole) fusion (*Baldehaar et al., 2010; Epp et al., 2011; Baldehaar et al., 2013; Karim et al., 2017*). Knocking out GYP6 partially suppresses protein trafficking defects observed in *nhx1*∆ cells, suggesting that in the absence of NHX1, Gyp6 may inactivate these Rab-GTPases preventing MVB membrane fusion events (*Ali et al., 2004*). Since this discovery, Fratti and colleagues reported that deleting NHX1 partially blocks homotypic vacuole fusion, and Ypt7––the Rab-GTPases responsible for this event––is not likely targeted (*Qiu and Fratti, 2010; Wickner, 2010*). However, this does not explain how Nhx1 contributes to endocytosis or MVB biogenesis, as Nhx1 is not present on vacuole membranes within living cells (*Nass and Rao, 1998; Kojima et al., 2012*), which is why it has been used as a reference protein to label endosomes in *S. cerevisiae* (e.g. *Huh et al., 2003*). More importantly, this fusion event does not contribute to endocytosis. However, most of the machinery that underlies homotypic vacuole fusion is also thought to mediate MVB-vacuole fusion (*Nickerson et al., 2009; Kümmel and Ungermann, 2014; Karim et al., 2017*) implicating Nhx1 in this process.

Thus, when considering all published findings, we reasoned that Nhx1 may contribute to MVB-vacuole membrane fusion, the last step of the endocytic pathway. Recently, our group developed a cell-free assay to study the molecular mechanisms underlying this fusion event in *S. cerevisiae* (*Karim et al., 2017*). Here we use it to test this hypothesis and begin to characterize how Nhx1 contributes to this process.

## RESULTS AND DISCUSSION

### Ionic requirements for MVB-vacuole fusion suggest dependence on Nhx1

All endosomal NHEs including Nhx1 contribute to ion gradients across endosome or MVB perimeter membranes where they export a lumenal H^+^ ion (to regulate pH) in exchange for import of a cytoplasmic monovalent cation (to possibly regulate the osmotic gradient; *Nass and Rao, 1998; Bowers et al., 2000*). As ionic and osmotic gradients are important for membrane fusion events underlying exocytosis and endocytosis as well as homotypic vacuole fusion *(Heuser, 1989; Starai et al., 2005; Brett and Merz, 2008; Cang et al, 2015*), we reasoned that Nhx1 activity may contribute to MVB-vacuole fusion. If true, then this fusion event should be dependent on pH and monovalent cation gradients. To test this hypothesis, we conducted a series of cell-free MVB-vacuole fusion assays that involve reconstitution of β-lactamase activity upon lumenal content mixing (see *Karim et al., 2017*). Importantly, Nhx1-positive puncta, representing endosomes or MVBs, are present and found adjacent to vacuole membranes in organelle preparations freshly isolated from yeast cells expressing NHX1-GFP (Figure 1A), similar to its distribution in living cells (*Nass and Rao, 1998*). In addition, we observe the accumulation of Nhx1-GFP on membranes of large vacuoles over time under fusogenic conditions in vitro, suggesting that Nhx1-GFP is deposited on products of fusion between MVB perimeter and vacuole membranes. These important observations justify the use of our cell-free system to study the potential role of Nhx1 in MVB-vacuole fusion. Furthermore, they support the idea that MVB membranes undergo full fusion (i.e. completely merge) with vacuole membranes to deliver lumenal contents (i.e. intralumenal vesicles) to the vacuole.

**Figure 1.**
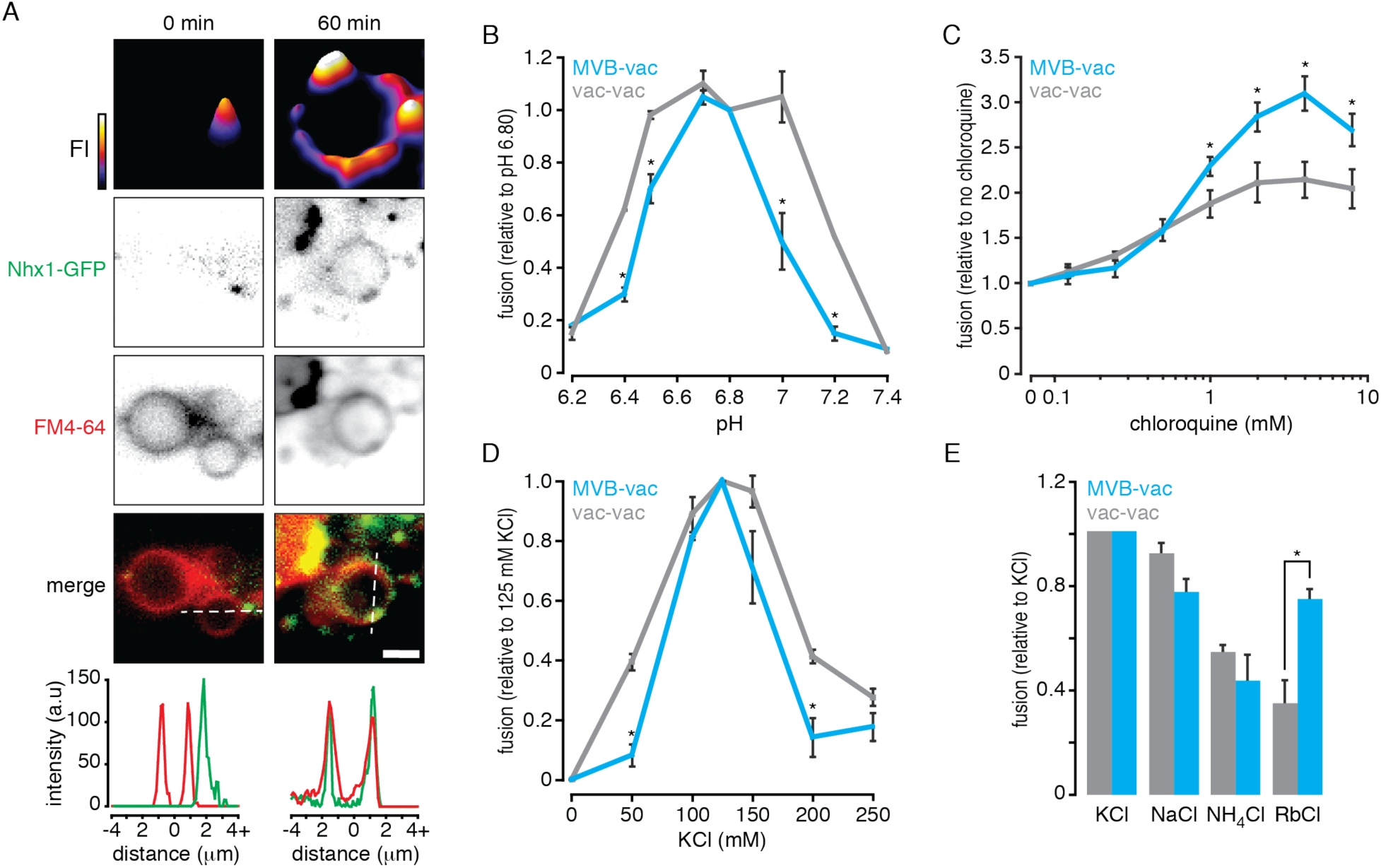
Ionic profile of MVB-vacuole fusion. (A) Fluorescence micrographs of organelles isolated from yeast cells expressing Nhx1-GFP before (0) and after (60 minutes) incubation under fusogenic conditions *in vitro*. Vacuole membranes are stained with FM4-64. 2-dimensional fluorescence intensity (FI) plots are shown for Nhx1-GFP (top). Fluorescence intensity is also plotted (bottom) for lines shown in merged images of Nhx1-GFP and FM4-64 channels. Scale bar, 2 µm. (B–E) MVB-vacuole (MVB-vac) or vacuole-vacuole (vac-vac) fusion measured in the presence of increasing pH (B), chloroquine (C), KCl (D), or when KCl was replaced with other salts (125 mM; E). All fusion reactions were incubated for 90 minutes at 27˚C in the presence of ATP. Values were normalized to control conditions (125 mM KCl, pH 6.80, no chloroquine). Mean ± S.E.M. (n ≥ 3) are plotted and P-values < 0.05 (*) are shown for comparisons between MVB-vacuole and vacuole-vacuole fusion values.

After confirming the presence of Nhx1 in our cell-free preparations, we next changed the pH of the reaction buffer to mimic changes in cytoplasmic [H^+^] and examined the effect on organelle fusion *in vitro* (Figure 1B). Compared to homotypic vacuole fusion, MVB-vacuole fusion was more sensitive to shifts in buffer pH and a sharp peak was observed at pH 6.80, the cytoplasmic pH of yeast cells under normal growth conditions (*Brett et al., 2005b; Mitsui et al., 2011*), suggesting that small changes in cytoplasmic pH are sufficient to control MVB-vacuole fusion events. We observed a different pH profile for homotypic vacuole fusion, suggesting that the mechanism(s) conferring pH-sensitivity are likely different for each fusion event.

In mammalian cells, alkalinization of the MVB lumen promotes fusion with lysosomes (*Cao et al., 2015*) consistent with our model of endosomal NHE function, whereby Nhx1 acts as a lumenal proton leak pathway opposing the activity of the V-type H^+^-ATPase (*Nass and Rao, 1998*). Thus, to determine if alkalinizing lumenal pH also affects MVB-vacuole fusion in our system, we treated isolated organelles with chloroquine, a weak base that accumulates within MVBs and vacuoles to raise lumenal pH *(Pearce et al., 1999; Qiu and Fratti, 2010*). As compared to homotypic vacuole fusion, chloroquine had a greater stimulatory effect on MVB-vacuole fusion (Figure 1C). This finding is consistent with observations in mammalian systems, as well as for homotypic vacuole fusion (*Desfougères et al., 2016*), and confirms that this heterotypic fusion event is particularly sensitive to changes in both lumenal and cytoplasmic pH, which are regulated by Nhx1 (*Brett et al., 2005b; Mitsui et al., 2011*).

In exchange for the export of H^+^, Nhx1 is thought to import a monovalent cation: K^+^, Na^+^ or Rb^+^, and other NHEs are also known to bind and transport NH_4_^+^ with less affinity (*Nass et al., 1997; Brett et al., 2005a; Brett et al., 2005b*). If Nhx1 activity is important for MVB-vacuole fusion, then this process should depend on the presence of these monovalent cations in the reaction buffer that mimics the cytoplasm in vitro. To test this hypothesis, we first measured organelle fusion in the presence of increasing concentrations of K^+^ (as KCl), the most abundant monovalent cation in the cytoplasm and likely preferred substrate of Nhx1 (Figure 1D). As expected, MVB-vacuole fusion required KCl and peak fusion was observed near physiological concentrations (125 mM). Heterotypic fusion was less tolerant to changes in [KCl] as compared to homotypic vacuole fusion, confirming that this fusion event is sensitive to both substrates of Nhx1.

We next tested if organelle fusion was supported by other cationic substrates by replacing K+ with Na+, Rb+ or NH4+ (Figure 1E). Replacing K+ with Na+ or Rb+ reduced heterotypic fusion by 20.4 and 22.4% respectively, whereas replacing it with NH4+ diminished heterotypic fusion by 41.2%, which is consistent with the predicted relative affinities of Nhx1 for each cation *(Nass and Rao, 1998; Brett et al., 2005a; Brett et al., 2005b*). Similar results were obtained for homotypic vacuole fusion – consistent with a previous report (*Starai et al., 2005*) – with the exception of Rb^+^, which does not entirely support homotypic fusion, possibly reflecting the absence of Nhx1 or other mechanisms capable of Rb^+^ transport on the vacuole. In all, these findings reveal the cationic profile of MVB-vacuole fusion, which is distinct from homotypic vacuole fusion and correlates with Nhx1 ion exchange activity.

### Deleting NHX1 impairs MVB-vacuole membrane fusion

Previous studies on Nhx1 function have led to the hypothesis that deleting NHX1 blocks MVB-vacuole fusion (*Bowers et al., 2000; Kallay et al., 2011*). To test this hypothesis, we knocked out NHX1 in cells expressing fusion probes and confirmed that GFP-tagged variants of the probes properly localize to the lumen of MVB or vacuole in live cells or organelles isolated from *nhx1*∆ strains (Figure 2A). Next, we measured fusion between MVBs and vacuoles isolated from either wild type or *nhx1*∆ cells (Figure 2B). As expected, deleting NHX1 impaired MVB-vacuole fusion, confirming that it contributes to this fusion event. However, knocking out NHX1 did not completely block this fusion event, consistent with Nhx1 possibly playing a regulatory role as opposed to being a component of the core fusion machinery which when deleted cause severe trafficking defects and vacuole fragmentation (i.e. *vps* class B or class C phenotypes) unlike *nhx1*∆ (a *vps* class E mutant; *Robinson et al., 1988; Rothman et al., 1989*).

**Figure 2.**
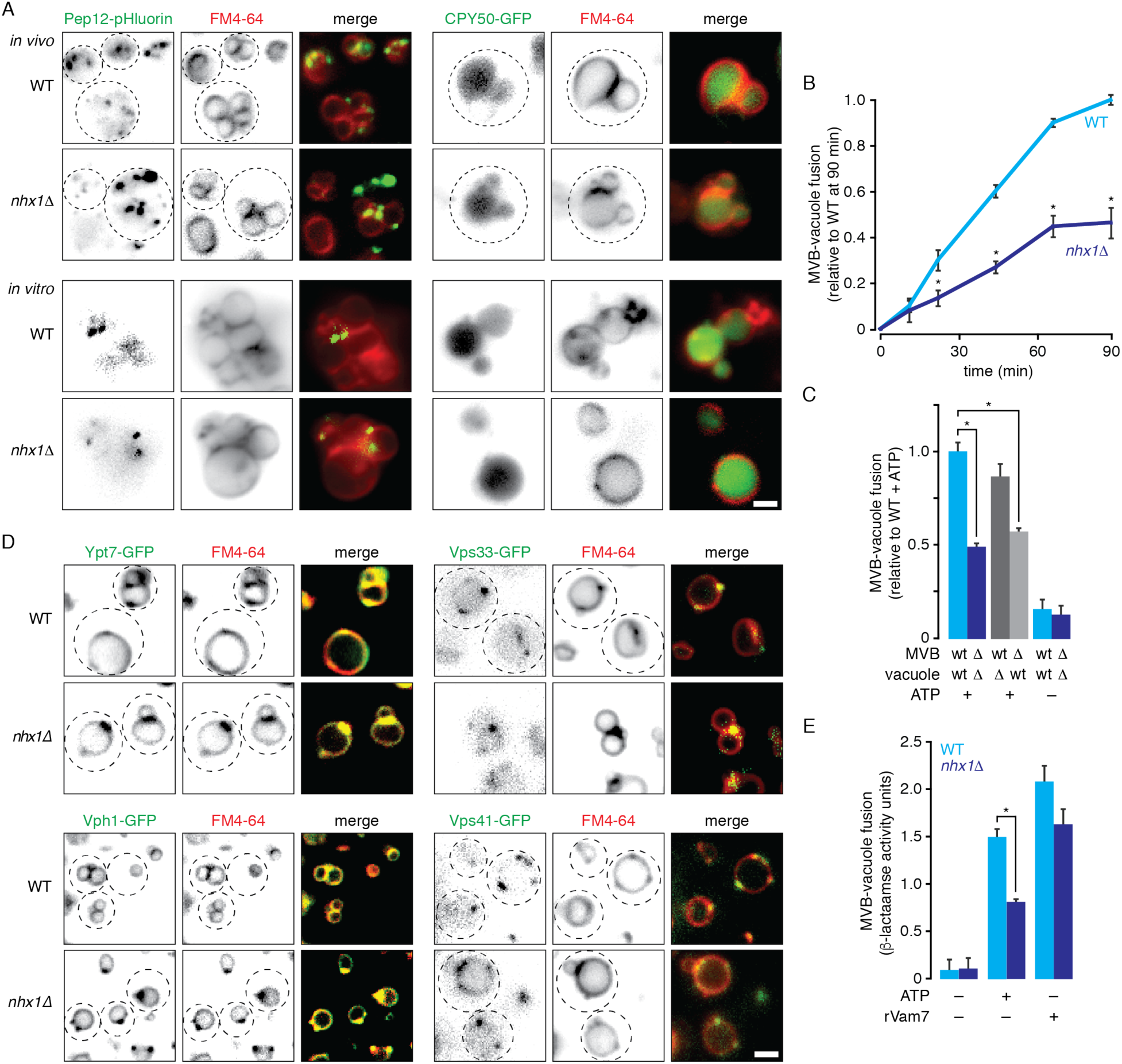
Deleting NHX1 impairs MVB-vacuole membrane fusion by targeting SNAREs. (A) Fluorescence micrographs of wild type (WT) or *nhx1Δ* cells (top) or isolated organelles (bottom) expressing Pep12-pHluorin or CPY50-GFP. Vacuole membranes are stained with FM4-64. (B) MVB-vacuole fusion between organelles isolated from WT or *nhx1*∆ cells measured over time in the presence of ATP. (C) MVB-vacuole fusion measured after mixing organelles from WT or *nhx1*∆ (∆) cells expressing either MVB or vacuole fusion probes. Reactions were incubated for 90 minutes at 27˚C in the absence or presence of ATP. (D) Fluorescence micrographs of WT or nhx1∆ cells expressing Ypt7, Vph1, Vps33 or Vps41 fused to GFP. Vacuole membranes are stained with FM4-64. (E) MVB-vacuole fusion between organelles isolated from WT or *nhx1*∆ cells in the presence or absence of ATP or 100 nM rVam7. Mean ± S.E.M. (n ≥ 3) are plotted and P-values < 0.05 (*) are shown for comparisons between wild type and *nhx1*∆ MVB-vacuole fusion values. Dotted lines outline cell perimeter as assessed by DIC. Scale bars, 2 μm.

Deleting NHX1 blocks delivery of some biosynthetic cargo to the vacuole (*Bowers et al., 2000; Brett et al., 2005b*). Thus, it is possible that impaired MVB-vacuole fusion may be a consequence of improper delivery of fusion proteins to vacuoles in *nhx1*∆ cells. To eliminate this possibility, we mixed organelles isolated from either wild type or *nhx1*∆ cells expressing complementary fusion probes and measured heterotypic membrane fusion *in vitro* (Figure 2C). Importantly, fusion between wild type MVBs and *nhx1*∆ vacuoles was similar to fusion between wild type organelles, confirming that the fusion machinery was properly delivered to vacuoles in *nhx1*∆ cells, consistent with a previous report (*Qiu and Fratti, 2010*). However, fusion between *nhx1*∆ MVBs and wild type vacuoles was impaired, similar to fusion between only *nhx1*∆ organelles. This important finding indicates that underlying fusion defect is inherent to the MVB membrane, where Nhx1 resides, not the vacuole membrane.

What prevents the MVB perimeter membrane from fusing with the vacuole membrane when NHX1 is deleted? One possibility is that the underlying fusion machinery, such as the Rab-GTPase Ypt7, its cognate multisubunit tethering complex HOPS (HOmotypic fusion and vacuole Protein Sorting complex) or SNAREs may be missing on MVB membranes in *nhx1*∆ cells. To test this hypothesis, we tagged Ypt7 or subunits of HOPS (Vps41 or Vps33) with GFP and examined their distribution within live yeast cells (Figure 2D). Ypt7-GFP, Vps41-GFP and Vps33-GFP were present on puncta (representing MVBs) and vacuole membranes in *nhx1*∆ cells, similar to their distributions in wild type cells (also see *Auffarth et al., 2014*). This finding is consistent with a previous work showing that these and other fusogenic proteins required for this fusion event (e.g. the SNAREs Vam3, Vam7 and Nyv1) are present at similar levels as wild type organelle preparations containing MVBs and vacuoles isolated from *nhx1*∆ cells (*Qiu and Fratti, 2010*). To confirm that SNAREs were functional on MVBs isolated from *nhx1*∆ cells, we next stimulated fusion in vitro with the soluble Qc-SNARE rVam7 in place of ATP, which drives trans-SNARE pairing and zippering to stimulate MVB-vacuole fusion, independent of Ypt7 and HOPS function (*Throngren et al., 2004*). Indeed, rVam7 rescued fusion of organelles isolated from *nhx1*∆ cells (Figure 2E). In all, these findings confirm that core fusion machinery is intact on MVBs isolated from *nhx1*∆ cells.

### Nhx1 does not target Ypt7 to regulate MVB-vacuole fusion

How does ion transport by Nhx1 affect the fusion machinery? It was shown previously that Nhx1 binds and inhibits Gyp6, a Rab-GTPase Activating Protein (Rab-GAP) that can inactivate Ypt7, the Rab-GTPase responsible for MVB-vacuole fusion *(Vollmer et al., 1999; Will and Gallwitz, 2001; Ali et al., 2004; Brett et al., 2008; Karim et al., 2017*). Thus, we hypothesized that in the absence of NHX1, Gyp6 should be stimulated and inactivate Ypt7 on MVB membranes to prevent MVB-vacuole fusion. To test this hypothesis, we first assessed the proportion of active Ypt7 protein present on isolated organelles by adding a purified, recombinant Gdi1, a Rab-chaperone protein (rGdi1) to isolated organelles. Gdi1 selectively binds and extracts inactive Rab from isolated membranes, allowing us to separate the pool of soluble rGdi1-bound inactive Ypt7 from membrane bound (presumably active) Ypt7 by differential centrifugation (*Brett et al., 2008*). Compared to organelles isolated from wild type cells, MVBs and lysosomes from *nhx1*∆ cells contained similar amounts of active Ypt7 on membranes (Figure 3A), and this pool of Ypt7 was equally susceptible to inactivation by Rab-GAP activity (by adding recombinant Gyp1-46 protein, which contains the catalytic TBC domain of the Rab-GAP Gyp1) or activation by addition of GTPγS a non-hydrolysable analog of GTP, suggesting that deleting NHX1 has no effect on Ypt7 activity.

**Figure 3.**
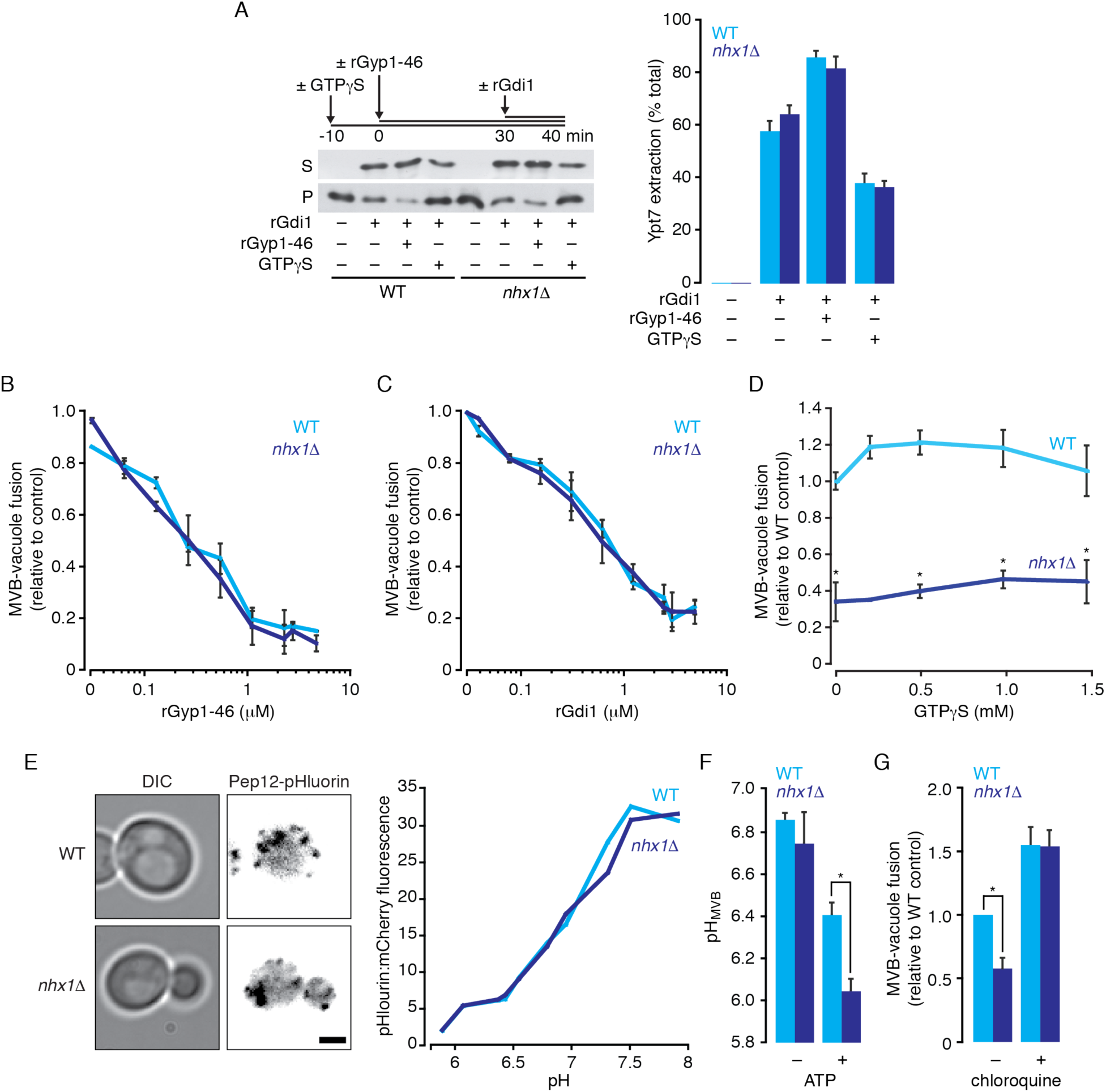
MVB-vacuole fusion defect caused by *nhx1*∆ is due to lumenal hyper-acidification, not Rab-GTPase inactivation. (A) Organelles isolated from wild type (WT) or *nhx1Δ* cells were incubated with ATP for 40 minutes with or without 10 µM rGdi1 during the last 10 minutes. Reactions were treated with 0.2 mM GTPγS or 5 µM rGyp1-46 where indicated. After incubation, membrane-bound (pellet, P) and soluble (supernatant, S) proteins were separated by centrifugation, and Ypt7 in each fraction was assessed by immunoblotting (left). Densitometric analysis of Ypt7 extraction (% total protein found in soluble fraction) is shown (right). (B-D) MVB-vacuole fusion between organelles isolated from WT or *nhx1*∆ cells measured in the absence or presence of increasing rGyp1-46 (B), rGdi1 (C) or GTPγS (D). Values were normalized to control conditions (no inhibitors or GTPγS). (E) pHluorin fluorescence and DIC micrographs of WT or *nhx1Δ* cells expressing Pep12-pHluorin-mCherry (left). Scale bar, 2 μm. (right) Calibration curves showing probe florescence at increasing pH values in both strains. (F) Lumenal pH of isolated MVBs from WT or *nhx1*∆ cells measured in the presence or absence ATP. (G) MVB-vacuole fusion between organelles isolated from WT or *nhx1*∆ cells measured in the absence or presence of 4 mM chloroquine. All fusion reactions were incubated for 90 minutes at 27˚C in the presence of ATP. Mean ± S.E.M. (n ≥ 3) are plotted and P-^values < 0.05 (*) are shown for comparisons between wild type and *nhx1*∆ MVB-vacuole fusion or pHMVB^ values.

To further test this hypothesis, we treated fusion reactions containing organelles isolated from WT or *nhx1*Δ cells with increasing concentrations of the Ypt7 inhibitors rGdi1 or rGyp1-46 (*Eitzen et al., 2000; Brett and Merz, 2008*). We reasoned that if less active Ypt7 was present on membranes in absence of NHX1, then MVB-vacuole fusion should be more susceptible to inhibition by these proteins. However, responses to rGdi1 or rGyp1-46 were similar in the presence or absence of NHX1 (Figure 3B and C), suggesting that loss of NHX1 does not affect the activation state of Ypt7.

These results are in stark contrast to the effect of deleting components of the ESCRT machinery (e.g. Vps27, a component of ESCRT-I; *Henne et al., 2011*), whereby abolishing ESCRT-mediated intralumenal vesicle formation prevents proper maturation of MVBs, which in turn, blocks subsequent fusion with vacuoles presumably by disrupting a Rab conversion mechanism required for Ypt7 activation (*Russell et al., 2012; Karim et al., 2017*). In support of this model, we have previously shown that replacing wild type YPT7 with a mutant locked in its active state (Ypt7-Q68L) rescues this fusion defect, as does addition of GTPγS to fusion reactions in vitro (which irreversibly activates Ypt7; *Karim et al., 2017*). Although contentious (see *Kallay et al., 2011*), Kanazawa and colleagues have proposed that knocking out NHX1 may prevent recruitment of Vps27 to endosome membranes, partially blocking ILV formation (*Mitsui et al., 2011*). If true, then defective MVB maturation caused by deleting NHX1 should also prevent Rab conversion and MVB-vacuole fusion should be rescued by GTPγS. However, when we tested this hypothesis, we found that addition of GTPγS did not rescue fusion between MVBs and vacuoles isolated from *nhx1*∆ cells (Figure 3D), consistent with results presented herein and previous reports (*Kallay et al., 2011*). Thus, in all, these results suggest that Nhx1 activity does not affect Ypt7 function, unlike perturbing ESCRT function, but rather targets another mechanism needed to promote efficient trans-SNARE pairing and zippering for MVB-vacuole fusion.

### Lumenal hyper-acidification correlates with a MVB-vacuole fusion defect in nhx1∆ cells

Deleting NHX1 hyper-acidifies the lumen of MVBs (or endosomes) and vacuoles in living cells and treating them with weak bases, such as chloroquine or methylamine, rescues endocytic trafficking defects (*Ali et al., 2004; Brett et al., 2005b; Mitsui et al., 2011)*. Specifically, they prevent accumulation of internalized surface proteins at the MVB within *nhx1*∆ cells and allow proper delivery to the vacuole lumen where they are degraded. This observation suggests that MVB-vacuole fusion defects caused by deleting NHX1 can be overcome by the addition of weak bases. However, prior to testing this hypothesis, we first confirmed that MVBs isolated from *nhx1*∆ were indeed hyper-acidic. Using a similar approach as that reported by Hiroshi Kanazawa and colleagues (*Mitsui et al., 2011*), we measured lumenal LE pH by tagging the lumenal face of endosomal Qa-SNARE Pep12 with a pH-sensitive variant of GFP called pHluorin. After demonstrating that the probe was properly localized to MVBs and responsive to pH in our organelle preparations (Figure 3E), we confirmed that enlarged MVBs isolated from *nhx1*∆ cells were hyper-acidified (Figure 3F), validating our use of this cell-free assay to study the role of Nhx1 in MVB-vacuole fusion. Importantly, we show that MVB acidification is ATP-dependent, confirming that ATP is needed to drive H^+^-pumping by the V-type H^+^-ATPase (or V-ATPase), in support the prevailing model of MVB pH regulation that describes Nhx1 as a lumenal H^+^ leak pathway that opposes V-type H^+^-ATPase function (*Nass and Rao, 1998*).

We next determined if addition of the weak based chloroquine rescues MVB-vacuole fusion defects caused by NHX1 deletion (Figure 3G). As predicted, chloroquine completely rescued MVB-vacuole fusion, consistent with previous reports that H^+^ transport by Nhx1 is essential for delivery of internalized surface membrane proteins to vacuoles for degradation in live cells (*Brett et al., 2005b*). Weak bases also rescue homotypic vacuole fusion defects caused by deleting NHX1 (*Qiu and Fratti, 2010*), suggesting the underlying mechanism may be similar.

What senses and responds to pH at MVBs to initiate fusion with vacuoles? Before we speculate, it is important to acknowledge that Nhx1 is thought to translocate H^+^ from the lumen to the cytoplasm; thus, alterations in pH on either side of the MVB perimeter membrane may contribute to the fusion defect caused by deleting NHX1. On the cytoplasmic side, loss of Nhx1 activity may deplete H^+^ near the outer leaflet of the MVB lipid bilayer. Data presented in Figure 1B suggests that even a small decrease in cytoplasmic [H^+^] (from pH 6.8 to 7.2) completely blocks fusion. MVB and vacuole membrane outer leaflets are enriched with negatively charged phospholipid species, e.g. phosphatidylinositide-3-phosphate and phosphatidic acid (*Gillooly et al., 2000; Fratti et al., 2004*). Many components or regulators of the fusion protein machinery are recruited to the membrane by binding these lipids, including the Ypt7 guanine exchange factor complex Mon1-Ccz1 (*Lawrence et al., 2014*), HOPS (*Stroupe et al., 2006*), Vac1 and Vps45 mediated by FYVE domains (*Peterson et al., 1999; Tall et al., 1999)*, and Vam7 through its PX domain (*Fratti and Wickner, 2007*). As it has been proposed that local H^+^ can protonate these lipids to modulate recruitment of the fusion protein machinery to the membrane (*He et al., 2009; Shin and Loewen, 2011; Starr et al., 2016*), it is possible that deleting NHX1 perturbs lipid protonation on the cytoplasmic leaflet disrupting recruitment of fusion proteins to the MVB membrane. Although appealing, we argue that this possibility is unlikely because: deleting NHX1 has no effect on pH near the cytoplasmic leaflet of the MVB membrane (*Mitsui et al., 2011*). Furthermore, we do not observe changes in cellular distributions of GFP-labeled components of HOPS (Figure 2D) and Ypt7 can be activated in *nhx1*∆ cells indicating that negatively charged lipids on MVB membranes continue to recruit soluble components of the fusion machinery.

Rather, we propose that the reduction in pH on the lumenal side of the MVB membrane contributes to the fusion defect in *nhx1*∆ cells. This is supported by two important observations: fusion is rescued by adding a weak base to raise lumenal pH (Figure 3G) or by adding rVam7 to stimulate fusion in the absence of ATP, a condition that does not allow lumenal hyper-acidification by the V-type H^+^-ATPase in vitro (see Figure 3F). However, the molecular basis of sensing lumenal pH within organelles of the endocytic system largely remains an enigma. Although, it has been suggested that subunits of the V-type H^+^-ATPase may sense lumenal H^+^ ion concentrations within the vacuole (or lysosome) to regulate fusion and other cellular roles (e.g. trigger cytoplasmic TOR signaling in response to cellular metabolism; *Maxson and Grinstein, 2014; Lim and Zoncu, 2016*). Although contentious (*Coonrod et al., 2013*), the V-type H^+^-ATPase is also implicated in the homotypic vacuole fusion reaction, whereby it was proposed to mediate pore formation, a critical intermediate of the lipid bilayer fusion (*Peters et al., 2001*). Interestingly, Vph1, the stalk domain of the V-ATPase that is exclusively found on vacuole membranes in wild type cells, is redistributed onto enlarged MVBs when NHX1 is deleted (*Bowers et al., 2000*). Thus, it is possible that the V-type H^+^-ATPase, possibly through its interactions with SNARE proteins (*Strasser et al., 2011*), may link Nhx1 function to MVB-vacuole fusion.

### Relevance to human disease

Mutations in the human orthologs of yeast NHX1, the endosomal Na^+^(K^+^) exchangers NHE6 and NHE9 (*Brett et al., 2002; Brett et al., 2005a; Hill et al., 2006*), are linked to neurological diseases such as Christianson syndrome, autism, attention deficit and hyperactivity disorder and epilepsy (*Gilfillan et al., 2008; Lasky-Su et al., 2008; Morrow et al., 2008; Markunas et al., 2010; Cardon et al., 2016; Yang et al., 2016*). However, the etiology is not entirely understood. Given that human NHE6 or NHE9 replace some functions of Nhx1 in *S. cerevisiae* (*Hill et al., 2006*) and that the genes encoding the machinery responsible for MVB-vacuole (or lysosome) fusion are found in all eukaryotes (*Luzio et al., 2010*), we predict that the cellular roles of endosomal NHEs are evolutionarily conserved. Indeed, compelling work by the Morrow, Walkley, Strømme and Orlowski groups demonstrates a strikingly similar role for mammalian NHE6 within neurons where it seems important for endocytosis of surface TrkB/Brain Derived Neurotropic Factor receptors needed for cell signaling events underlying proper development of neural circuitry (*Strømme et al., 2011; Ouyang et al., 2013; Deane et al., 2013*). Mutations in NHE6 or NHE9 are also thought to impair endocytosis in astrocytes possibly underlying cellular defects that contribute to Alzheimer’s disease (*Prasad and Rao, 2015*) or glioblastoma (*Kondapalli et al., 2015*). In all cases, impairment of endosomal NHE function correlates with hyper-acidification of endosomes (or MVBs; *Kondapalli et al., 2013; Ouyang et al., 2013*). Thus, it is tempting to speculate that loss-of-function mutations in NHE6 or NHE9 also disrupt MVB-lysosome membrane fusion to impair endocytosis contributing to the pathogenesis of these human diseases.

## MATERIALS AND METHODS

### Yeast strains and reagents

For organelle content mixing assays, we used the *Saccharomyces cerevisiae* strain BJ3505 [*MATα;pep4::HIS3;prbΔ1-1.6R;his3–Δ200;lys2–801;trp1-Δ101(gal3);ura3–52;gal2;can1*] transformed with expression plasmids containing the MVB fusion probe (pCB002 and pCB003) or vacuole fusion probe (pYJ406-Jun-Gs-α and pCB011; see *Karim et al., 2017*). NHX1::GFP was knocked in or NHX1 was knocked out of BJ3505 cells using the Longtine method (*Longtine et al., 1998*). To confirm proper fusion probe localization, WT or *nhx1*∆ BJ3505 cells were transformed with an expression plasmid containing the MVB targeting sequence (Pep12) fused to pHluorin (pCB035) or the vacuole targeting sequence (CPY50) fused to GFP (pCB044; see Karim et al., 2017). Reagents were purchased from Sigma-Aldrich, Invitrogen or BioShop Canada Inc. Purified rabbit polyclonal antibody against Sec17 or Ypt7 were gifts from William Wickner (Dartmouth College Hanover, NH, USA) and Alexey Merz (University of Washington, Seattle, WA, USA), respectively. Recombinant Gdi1 (*Brett et al., 2008*), Gyp1-46 (the catalytic domain of the Rab-GTPase activating protein Gyp1; *Eitzen et al., 2000*), Vam7 (*Schwartz and Merz, 2009*), or c-Fos (*Jun and Wickner, 2007*) proteins were purified as previously described. Reagents used in fusion reactions were prepared in 10 mM Pipes-KOH, pH 6.8, and 200 mM sorbitol (Pipes-sorbitol buffer, PS).

### Organelle isolation and membrane fusion

Organelles were isolated from yeast cells by the ficoll method as previously described (*Karim et al., 2017*). To assess organelle membrane fusion, organelle content mixing was measured using a complementary split β-lactamase based assay (see *Karim et al., 2017; Jun and Wickner, 2007*). In brief, organelles were isolated from separate yeast strains expressing a single fusion probe targeting either the MVB or vacuole. 6 μg of organelles isolated from each complementary strain were added to 60 μl fusion reactions in standard fusion buffer (125 mM KCl, 5 mM MgCl2, 10 μM CoA in PS) supplemented with 11 μM recombinant c-Fos protein to reduce background caused by lysis. ATP regenerating system (1 mM ATP, 40 mM creatine phosphate, 0.5 mg/ml creatine kinase) or 100 nM recombinant Vam7 protein (and 10 μg/ml bovine serum albumin) were added to stimulate fusion. Reactions were incubated up to 90 minutes at 27°C and then stopped by placing them on ice. Content mixing was quantified by measuring the rate of nitrocefin hydrolysis by reconstituted β-lactamase. 58 µl of the fusion reaction were transferred into a black 96-well clear-bottom plate and mixed with 142 µl of nitrocefin developing buffer (100 mM NaPi pH 7.0, 150 µM nitrocefin, 0.2% Triton X-100). To measure nitrocefin hydrolysis, absorbance at 492 nm was monitored at 15 seconds intervals for 15 minutes at 30°C with a Synergy H1 plate reader, a multimode spectrophotometer (Biotek, Winooski, VT, USA). Slopes were calculated and one fusion unit is defined as 1 nmol of hydrolyzed nitrocefin per minute from 12 µg of organelle proteins. Where indicated, purified antibodies raised against Sec17, purified recombinant Gdi1 or Gyp1-46, GTPγS, chloroquine were added to fusion reactions at concentrations indicated. In addition, pH or [KCl] of the fusion buffer was adjusted or KCl was replaced with equimolar (125 mM) NaCl, NH4Cl or RbCl as indicated. Experimental results shown are calculated from ≥ 3 biological replicates, each repeated twice (≥ 6 technical replicates total).

### Ypt7 extraction assay

Ypt7 extraction assays were performed as previously described (*Brett et al., 2008*). Briefly, 5X reactions containing 30 μg organelles isolated from WT or *nhx1Δ* cells either were incubated at 27°C for 30 minutes under fusion conditions. As a positive control, 5 µM rGyp1-46 was added to promote Ypt7 inactivation and extraction; as a negative control, organelles were pretreated with 0.2 mM GTPγS prior for 10 minutes prior to incubation to promote Ypt7 activation preventing extraction. 10 μM rGdi1 was then added and reactions were further incubated for 10 minutes. Samples were then immediately centrifuged (5,000 *g*, 5 minutes, 4°C) to separate membrane-bound (pellet) from soluble (supernatant) Ypt7 protein. Pellets were resuspended in 100 μl 1X of reaction buffer and 100 μl of the supernatant were mixed with 25 μl 5X Laemmli sample buffer, and then boiled for 10 minutes. One tenth of the total fraction volume was analyzed by immunoblotting using a purified rabbit antibody-raised against Ypt7 (see *Karim et al., 2017*). Western blots shown are best representatives of 3 biological replicates, each repeated twice (6 technical replicates total).

### MVB pH measurement

Based on a previous method used to measure lumenal MVB pH (*Mitsui et al., 2011*), we transformed wild type or *nhx1Δ* BJ3505 cells with pCB046, a yeast expression vector containing a genetically encoded pH probe (pHlourin fused to mCherry) fused to the C-terminus of Pep12 to exclusively target it to the lumen of MVBs. To generate calibration curves shown in Figure 3E, 12 μg of organelles were added to 60 μl reaction buffer containing 50 µM nigericin in PS buffer titrated to pH values between 5.8 and 8.0, and incubated at 27°C for 10 minutes. Fluorescence intensities for pHluorin (excitation at 485 nm, emission at 520 nm) and mCherry (excitation at 584 nm, emission at 610 nm) were then measured with a Synergy H1 plate-reader, a multimode spectrophotometer (Biotek, Winooski, VT, USA). A blank reference well containing 60 µl PS was used to detect background fluorescence. Ratios of background subtracted pHluorin:mCherry fluorescence values were then plotted versus pH. Fluorescence values obtained under standard fusion conditions (Figure 3F) were then compared to this curve to estimate MVB pH in vitro. Experimental results shown are best representatives of 3 biological replicates, each repeated twice (6 technical replicates total).

### Fluorescence microscopy

Live yeast cells were stained with FM4-64 to label vacuole membranes using a pulse-chase method as previously described (*Brett et al., 2008*). Membranes of isolated organelles were stained by adding 3 μM FM4-64 and incubating at 27°C for 10 minutes. Stained organelles were then added to standard fusion reaction buffer, incubated at 27°C for up to 60 minutes, and placed on ice prior to visualization. To obtain fluorescence micrographs, we used a Nikon Eclipse TiE inverted microscope equipped with a motorized laser TIRF illumination unit, Photometrics Evolve 512 EM-CCD camera, an ApoTIRF 1.49 NA 100x objective lens, and bright (50 mW) blue and green solid-state lasers operated with Nikon Elements software (housed in the Centre for Microscopy and Cellular Imaging at Concordia University). Micrographs were processed using ImageJ software (National Institutes of Health) and Adobe Photoshop CC. Images shown were adjusted for brightness and contrast, inverted and sharpened with an unsharp masking filter. Micrographs shown are best representatives of at least 3 biological replicates, imaged at least 6 times each (a total of ≥ 18 technical replicates) whereby each field examined contained ≥ 35 cells or isolated organelles.

### Data analysis and presentation

All quantitative data was processed using Microsoft Excel v.14.0.2 software (Microsoft Cooperation, Redmond, WA, USA), including calculation of means, S.E.M.s and student two-tailed t-tests. P-values < 0.05 are indicated. Fusion values shown were normalized to values obtained under control conditions (organelles isolated from wild type cells; pH 6.80; 125 mM KCl; ATP; no inhibitors, chloroquine or GTPγS) where indicated. Data was plotted using Kaleida Graph v.4.0 software (Synergy Software, Reading, PA, USA). All figures were prepared using Adobe Illustrator CC software (Adobe Systems, San Jose, CA, USA). References were prepared using Mendeley software (Mendeley, New York, NY, USA).

## AUTHOR CONTRIBUTIONS

M.A.K. performed all experiments and prepared all data for publication. M.A.K. and C.L.B. conceived the project and wrote the paper.

## ACKNOWLEDGEMENTS

We thank A.J. Merz and W.T. Wickner for antibodies. This work was supported by NSERC grants RGPIN/403537-2011 and RGPIN/2017-06652 awarded to C.L.B.

## LIST OF ABBREVIATIONS

ESCRT: Endosomal Sorting Complexes Required for Transport
GAP: Gtpase Activating Protein
HOPS: HOmotypic fusion and vacuole Protein Sorting
MVB: MultiVesicular Body
NHE: Na^+^/H^+^ Exchanger
SNARE: SNAp REceptor
VPS: Vacuole Protein Sorting
WT: Wild Type

